# Tools for genetic mapping and gene identification in *Caenorhabditis briggsae*

**DOI:** 10.64898/2026.11.05.764684

**Authors:** Haimeng Lyu, Nikita S. Jhaveri, Lance M. O’Connor, Robyn E. Tanny, Katie Freytag, Carli M. Genova, Bhagwati Gupta, Erik C. Andersen, Helen M. Chamberlin

## Abstract

The nematode *Caenorhabditis briggsae* serves as an important companion species for *Caenorhabditis elegans* for comparative studies. Because *C. briggsae* exhibits androdioecious reproduction and ease of laboratory culture similar to *C. elegans*, it has long been used for genetic studies in cross-species comparisons of mutant phenotypes to uncover the evolution of gene function. Here, we provide an updated toolkit for *C. briggsae* genetic experiments, leveraging the resource of recently improved *C. briggsae* genome sequence assemblies with variant mapping tools and gene identification methods. We identify and validate a set of indel variants between CGC2 (AF16) and two divergent strains, HK104 and VX34. We then use these variants to genetically map 10 previously isolated mutations and identify the impacted genes using high-throughput sequencing. We combine these new genes with others curated from the literature into an updated map with 73 genes, providing an enriched resource for *C. briggsae* genetic studies.

**Article Summary:** This project provides an updated toolkit for genetic studies in the nematode *Caenorhabditis briggsae*, an important comparison species for research on other *Caenorhabditis* species such as *Caenorhabditis elegans*. It combines the validation of indel variants with improved genome assembly data to produce an updated map and genetic methods useful to identify new mutations and to build strains with known mutations.

## Introduction

Comparative genomics provides a powerful approach to understanding genes, their functions, and their roles in biological processes, and comparative genomic methods are important in uncovering the differences responsible for the phenotypic diversity and innovations observed in individuals, populations, and species (Berner and Salzburger 2015; Broekema et al. 2020; Bomblies and Peichel 2022). However, evolution can alter the relative contributions and underlying biological networks even in the absence of overt phenotypic change. These differences are apparent only under system perturbation (such as environmental stress or mutation), but they fundamentally alter how the system works and how it can evolve (True and Haag 2001; Wagner 2005; Draghi et al. 2010; Zheng et al. 2019; Hulst et al. 2026). Consequently, a full understanding of the flexibility and constraints of a given genome, and how its functional characteristics are similar or distinct from others, relies on availability of both genetic and genomic tools that permit the discovery and characterization of mutations that alter gene function.

Nematodes of the *Caenorhabditis* genus are routinely used to investigate biological questions ranging across multiple scales (populations, organisms, cells) and uncover how genetic and genomic differences impact phenotypic outcomes (Baird et al. 2005; Hill et al. 2006; Félix 2007; Rockman and Kruglyak 2009; Gaertner and Phillips 2010; Ben-David et al. 2021). *Caenorhabditis elegans* is the best studied species of the genus, where research is facilitated by high-quality genomic resources, and a rich set of genetic mutations and tools (Ambros et al. 2025). Although several other species of the genus are routinely used for comparative studies, *Caenorhabditis briggsae* is distinct as an important comparison species and is one where genetic approaches, including genetic screens, generation of CRISPR alleles, and genetic mapping, have been pursued by many research groups (Hill et al. 2006; Inoue et al. 2007; Kelleher et al. 2008; Zhao et al. 2008; Baldi et al. 2009; Guo et al. 2009; Seetharaman et al. 2010; Zhao et al. 2010; Chamberlin et al. 2020; Velayudhan and Ellis 2022; Ivanova and Moss 2023). *C. briggsae* was the second species in the genus (after *C. elegans*) to have a complete draft genome sequence, greatly improving comparative genomic and genetic studies (Stein et al. 2003; Hillier et al. 2007). Since completion of the original work, advances in sequencing and genome assembly technologies have prompted the recent completion of updated, telomere-to-telomere genome assemblies and gene model curation for several *C. briggsae* strains, providing improved genomic resources for this important comparative species (Stevens et al. 2022; Moya et al. 2023; O’Connor et al. 2026).

*C. briggsae* mutants are routinely produced either using targeted genome-editing methods (Wei et al. 2014; Toker and Hobert 2022; Ivanova and Moss 2023; Lyu et al. 2024) or using forward genetic screening methods (Kelleher et al. 2008; Guo et al. 2009; Seetharaman et al. 2010; Sharanya et al. 2012; Sharanya et al. 2015; Chamberlin et al. 2020) for comparative studies. Although targeting methods permit precise engineering and the production of alleles directly orthologous to those identified in *C. elegans*, genetic screens remain important because of their potential for discovery, identifying genes or alleles that confer novel phenotypes or that uncover species-specific gene functions. Mapping by sequencing approaches, in which high-throughput sequencing methods are applied to pools of DNA from mutant and non-mutant F2 animals from F1 animals heterozygous for the mutation of interest (in one strain background) and the wild type (from a strain with a high degree of sequence variants compared to the first strain), are commonly used for *C. elegans* and other organisms, including *C. briggsae* (Doitsidou et al. 2016). Despite the power of these approaches, the availability of more classical mapping tools, such as visible genetic markers or defined polymorphic loci identifiable using PCR methods, is an important element of any genetic toolkit. They permit preliminary confirmation that a given trait is indeed genetically based, how many genes could be involved, and what the genetic properties are (Fay 2013). They are essential for construction of experimentally useful strains such as double mutants and provide resources for genetic balancing or tracking. Finally, from a teaching and training perspective, observing the mechanics of genetic transmission and recombination is often easier with a small set of loci before learners consider genome-scale data.

Here, we use the improved genomic resources to produce a suite of updated genetic tools for *C. briggsae*. We identify and validate DNA sequence variants among four *C. briggsae* strains, including marker sets identified between two genome assemblies from early *C. briggsae* studies (AF16 (now defined as CGC2) and HK104 (Koboldt et al. 2010; Ross et al. 2011)) and sets from two strains with improved genome assemblies and annotations (QX1410 and VX34 (Stevens et al. 2022)). We then apply these markers to the genetic mapping of 10 distinct mutations isolated by several research groups in the CGC2 genetic background, using a set of standardized protocols. We reference these genetic map data against whole-genome sequence data of the mutant strains, to identify the likely causative lesion and impacted gene for each strain. Finally, we integrate these genes into a curated map that incorporates *C. briggsae* genes identified by mutations for which the impacted gene is identified. Together, these results provide a robust updated toolkit for genetic studies in *C. briggsae*.

## Results

### Validation of a set of insertion-deletion (indel) variants across four wild-type C. briggsae strains using a standardized PCR-based protocol

Using strains bearing sequence variants compared to a standard reference strain has long been recognized as an important tool in comparative genomics, and the data from such strains are important in producing high-resolution genetic recombination maps (Ruvkun et al. 1989; Wicks et al. 2001). Genome sequence assemblies for reference *C. briggsae* strains (CGC2 and QX1410) included comparative analysis, variant calling, and production of genetic recombination maps using the divergent strains HK104 or VX34 (Koboldt et al. 2010; Stevens et al. 2022). Although previous studies have validated a set of single-nucleotide variant (SNV) and insertion-deletion (indel) loci between CGC2 and HK104 (Koboldt et al. 2010), we aimed to identify a set of indel variants across all four strains, using a common set of PCR conditions (Figure 1; Materials and Methods). We selected 35 sites with indels ranging from 7-32 bp from the CGC2/HK104 set (previously validated on CGC2 and HK104 DNA), and 72 sites with indels ranging from 50-100 bp from the QX1410/VX34 set, and used primers designed to observe the original variants on DNA templates from each of the four strains (CGC2, QX1410, HK104, VX34; Supplemental Tables 2, 3). Under a common set of PCR conditions, 32 (91%) of the CGC2/HK104 and 49 (68%) of the QX1410/VX34 set yielded robust products of the predicted size on DNA from the original two genome sequences used to predict the indels. Seventy of these 81 (86%) validated loci exhibited a similar product on both CGC2 and QX1410, consistent with the relatedness of these two strains (Stevens et al. 2022). For mapping mutations isolated in the CGC2 background, this analysis identified 40 validated polymorphic indels compared to HK104 (32 from the original CGC2/HK104 set plus 8 from the QX1410/VX34 predictions), and 59 compared to VX34 background (49 from the QX1410/VX34 set plus 10 from the CGC2/HK104 predictions), distributed across all six chromosomes (Supplemental Table 4).

**Figure 1.**
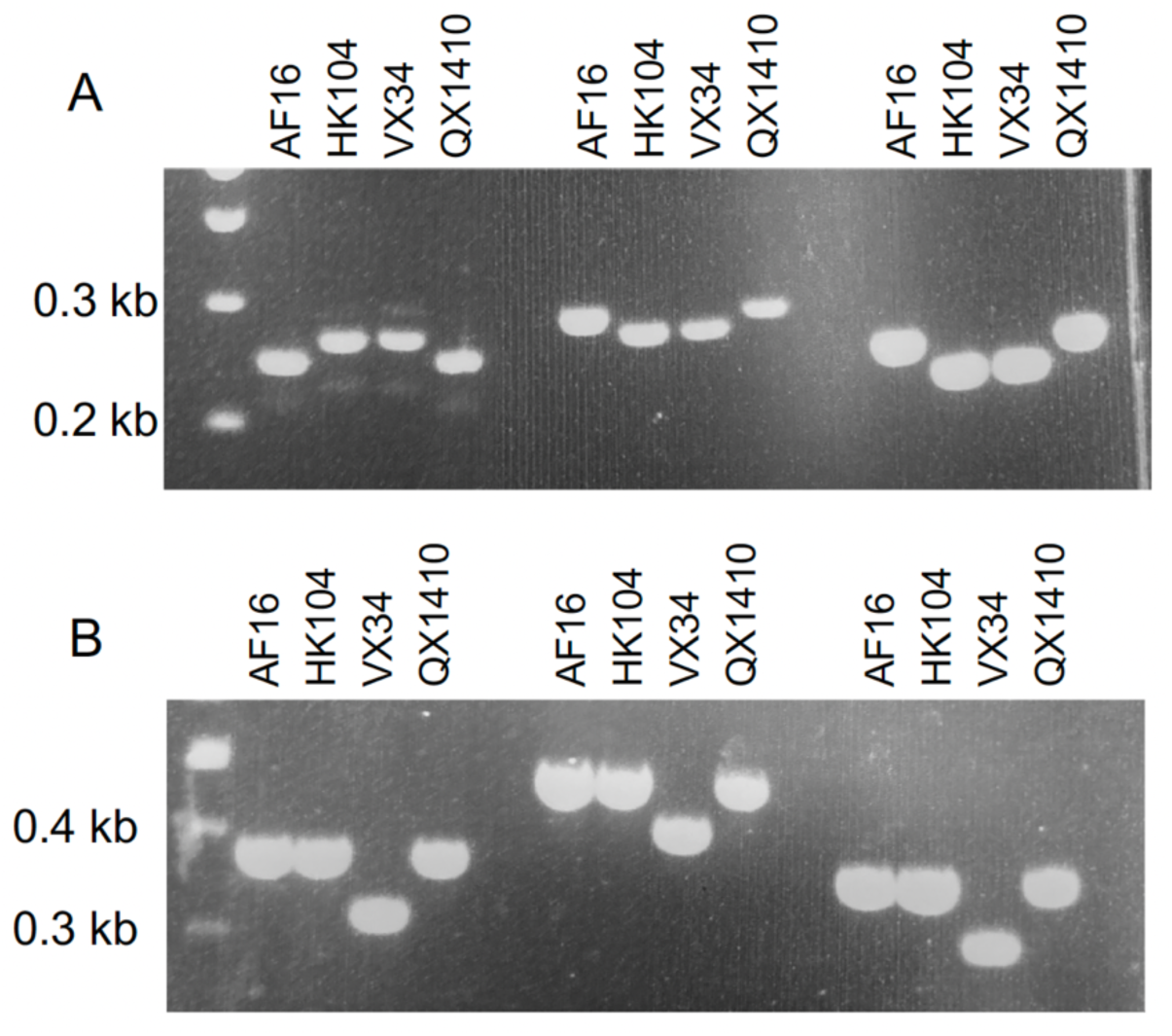
Representative agarose gel results for indel polymorphisms. Primers predicted to identify indels between CGC2 and HK104 from (Koboldt et al. 2010) (A) or between QX1410 and VX34 from Caendr.org (B). A. Left to right, PCR products amplified with primers flanking bhP9 (2325+2326), bhP5 (2331+2332), bhP26 (2337+2338). B. Left to right, PCR products amplified with primers flanking guP46 (6935+6936), guP48 (6937+6938), guP50 (6939+6941).

### Mutations isolated in CGC2 background can be mapped using HK104 or VX34

To demonstrate the use of these variants in genetic mapping experiments, we selected ten previously isolated *C. briggsae* mutants that exhibited defects in movement (uncoordinated (Unc)) or body morphology (shorter and/or smaller (dumpy (Dpy) or small (Sma)) (Supplemental Table 1). These mutations were isolated in standard F2 genetic screens, using EMS as a mutagen (Anderson 1995). The list included some genes with preliminary map data and even tentative gene assignments (such as *Cbr-unc-22(s1270),* based on its unique phenotype), as well as others that lacked detailed characterization and were isolated in a range of different laboratories. Although all mutations were nominally isolated in the CGC2 genetic background, selecting strains from different sources allowed us to evaluate the robustness of the marker loci for mapping in the context of potential genetic differences that could accumulate over time due to laboratory culture methods or exposure to mutagen. The mutation in five strains was mapped against both HK104 and VX34, and five were mapped solely against VX34 (Figure 2; Supplemental Tables 5, 6). Most of the mutations were initially assigned to a chromosome using a bulk-segregant method, wherein mutant and non-mutant F2 animals from parents heterozygous for the mutation (CGC2 background) and wild-type (HK104 or VX34 background) are selected into separate pools (Doitsidou et al. 2016). Alternatively, and for subsequent mapping on specific chromosomes, individual F2 homozygotes (displaying the recessive mutant phenotype or the wild-type phenotype for alleles that confer a dominant mutant phenotype) were selected individually to plates, and allowed to self, producing a DNA source that represents the product of two gametes from the heterozygous parent. Genotyping of these DNAs permit the calculation of recombination frequency between the polymorphic locus and the impacted gene, as well as defining clear genomic boundaries to limit the gene location (Supplemental Table 6). In these tests, all ten mutations behaved as single genes and mapped to a defined genomic region. The data for the five mutations mapped against both HK104 and VX34 and the defined regions were consistent between the two strains (Supplemental Tables 5, 6).

**Figure 2.**
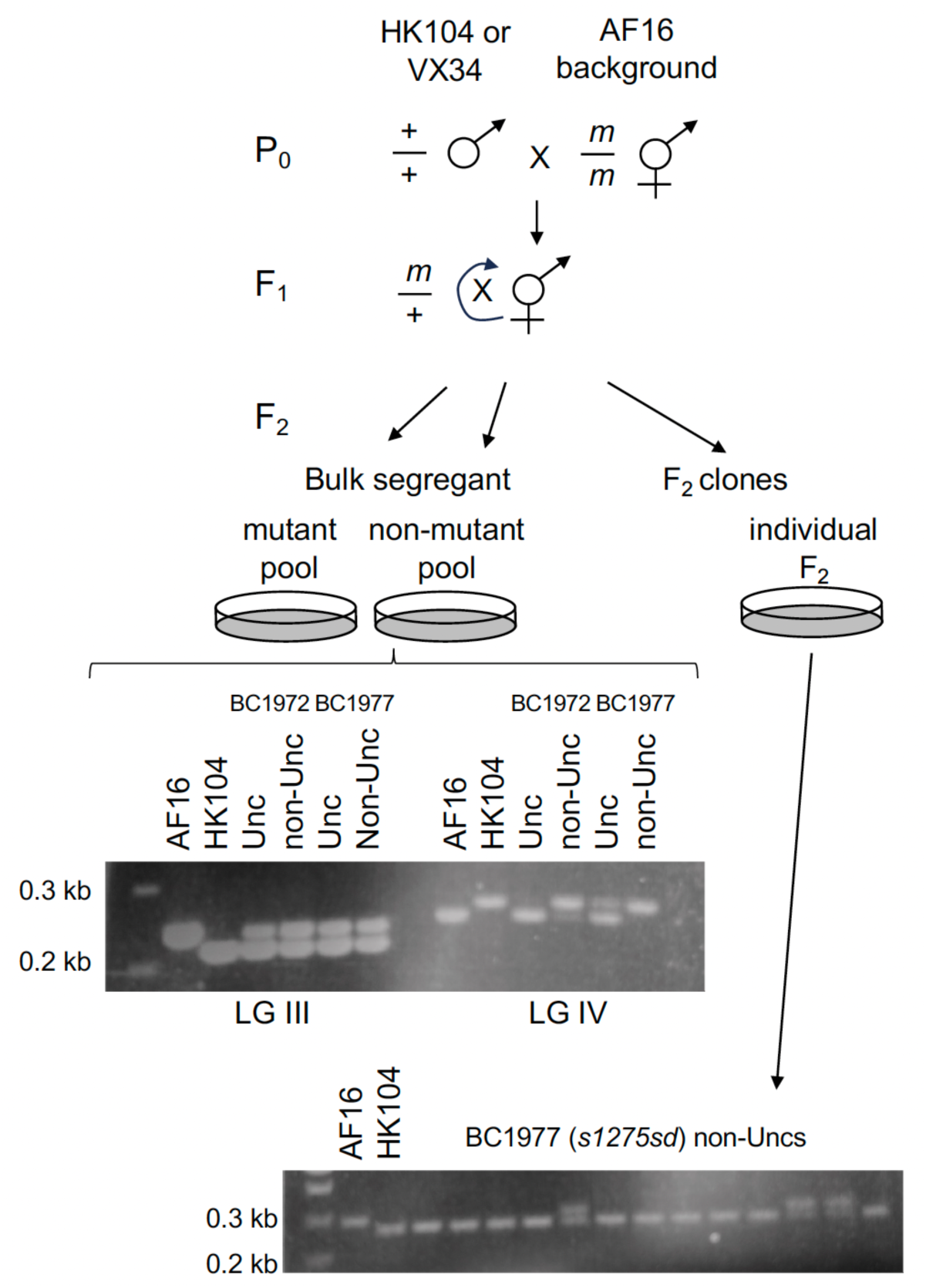
Crossing schemes for bulk-segregant and individual F2 mapping methods. Bulk-segregant representative data for BC1972 and BC1977. Products were amplified using primers flanking bhP14 (LGIII; 2319+2320) and flanking bhP9 (LGIV; 2325+2326). A sample of individual F2 tests for BC1977. Products were amplified using primers flanking cb-s214 (LGIV; 3401+3402). Although most F2 tests were carried out on plates of F2 mutant animals, the mutation in BC1977 causes a semi-dominant phenotype. Therefore, mapping was carried out by selecting against the mutation, cloning F2 wild-type (non-Unc) animals, with non-recombinants bearing the HK104 allele and recombinants bearing the CGC2 allele.

### Molecular identification of lesions associated with mutant strains links them to the physical map

To identify the affected genes and molecular lesions associated with each mapped mutation, we used two approaches. First, we used the genetic mapping and comparison to the *C. elegans* genome to identify likely candidate genes, with the assumption that the function and disrupted phenotype for this category of gene would be well conserved. This approach yielded three candidate genes (*Cbr-dpy-4* in CM3378, *Cbr-dpy-10* in CP4, *Cbr-dpy-6* in PB101) for which we used PCR amplification and Sanger sequencing to evaluate the candidate gene sequence in the mutant strains. This approach identified a premature stop codon that impacted each gene at, or earlier than, the position of nonsense or frame-shift alleles that similarly impact function of the orthologous gene in *C. elegans*, permitting assignment of the mutation to the gene (Figure 3, Supplemental Table 7, (Levy et al. 1993; Simmer et al. 2003)).

**Figure 3.**
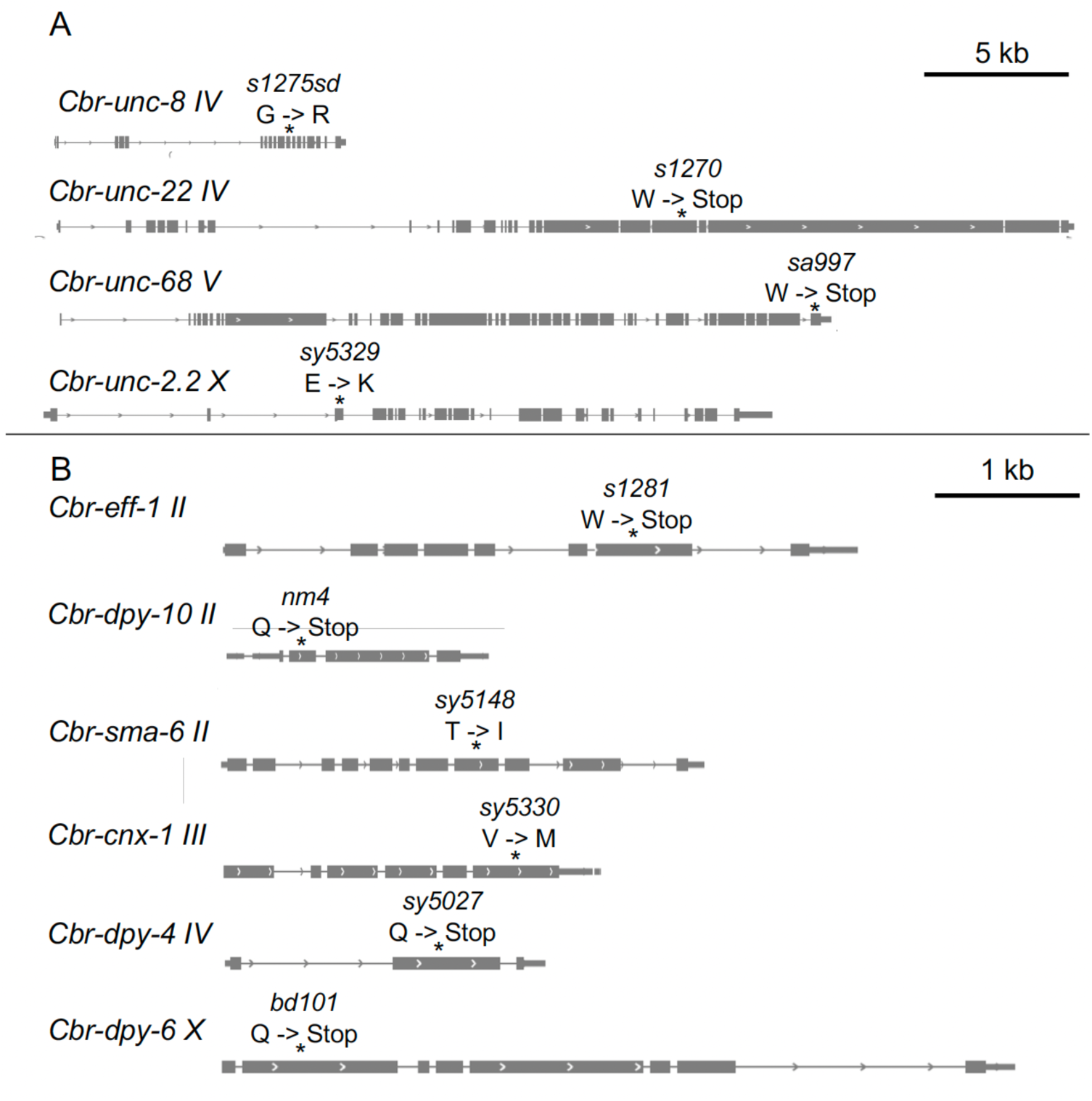
Genes with lesions identified in *C. briggsae* mutant strains. Gene model images adapted from CaeNDR.org (Crombie et al. 2024). A. Lesions identified in larger genes: *Cbr-unc-8* (BC1977), *Cbr-unc-22* (BC1972), *Cbr-unc-68* (JT10275), *Cbr-unc-2.2* (PS9334 Unc). B. Lesions identified in smaller genes: *Cbr-eff-1* (BC1983), *Cbr-dpy-10* (CP4), *Cbr-sma-6* (PS9148), *Cbr-cnx-1* (PS9334 Sma), *Cbr-dpy-4* (CM3378), *Cbr-dpy-6* (PB101).

For the remaining mutations, either a single candidate did not clearly emerge or the candidate gene was sufficiently large to make the single-gene sequencing approach untenable. Consequently, we sequenced the whole genomes of these mutant strains and then compared these sequences to the wild-type reference strain to identify EMS-derived mutations. The unique sequence variants for each strain were cross-referenced against the genetic mapping boundaries to define an interval where the candidate gene likely is found (Figure 3, Supplemental Tables 8-14). By evaluating the variants in each interval for their potential to impact gene function and the affected genes for orthologous function in *C. elegans*, we identified a single, strong candidate gene within each interval. Genes in two of the strains (*Cbr-unc-22* in BC1972; *Cbr-eff-1* in BC1983) met the same orthologous premature stop criteria as above (del Campo et al. 2005; Flibotte et al. 2010). JT10275 harbors a premature stop codon in the final exon of *Cbr-unc-68*. Although late in the coding sequence, this mutation is anticipated to truncate critical residues of the conserved Carboxyl-terminal domain (CTD) of the encoded Ryanodine Receptor (RYR, Figure 4A) that plays an important role in stabilization of the RYR channel pore during opening and closing of the channel, as well as interacting with channel activating ligands (Clarke and Hendrickson 2016; Peng et al. 2016). In BC1977, the *Cbr-unc-8* gene includes a G to R missense substitution that alters a residue conserved between Cbr-UNC-8 and Cel-UNC-8, in a domain common to DEG/ENaC family members shown to be important in channel closing (Figure 4B, (García-Añoveros et al. 1995)). Substitutions at this position are similarly associated with alleles of *Cel-unc-8* that confer a dominant phenotype (Tavernarakis et al. 1997). The mapped Unc phenotype in PS9334 coincides with a substitution (E to K) in Cbr-UNC-2.2 that impacts an amino acid immediately after the first transmembrane domain that is conserved in Calcium channel subunit alpha-1A proteins (Figure 4C, (Huang et al. 2019)). In PS9148, the *Cbr-sma-6* gene includes a T to I substitution, affecting a conserved residue in the kinase domain of BMP type 1 receptors, proximal to the conserved HRD motif of the catalytic loop (Figure 4D, (Gizzio et al. 2024)). The mapped Sma phenotype in PS9334 coincided with a V to M substitution in the *Cbr-cnx-1* gene. This residue, although conserved between Cel-CNX-1 and Cbr-CNX-1, is not in a domain conserved in Calnexin proteins from humans and flies (Lee et al. 2005). However, *Cel-cnx-1* null mutants are shorter than the wild type (Lee et al. 2005), consistent with the observed Sma phenotype of the *C. briggsae* mutant.

**Figure 4.**
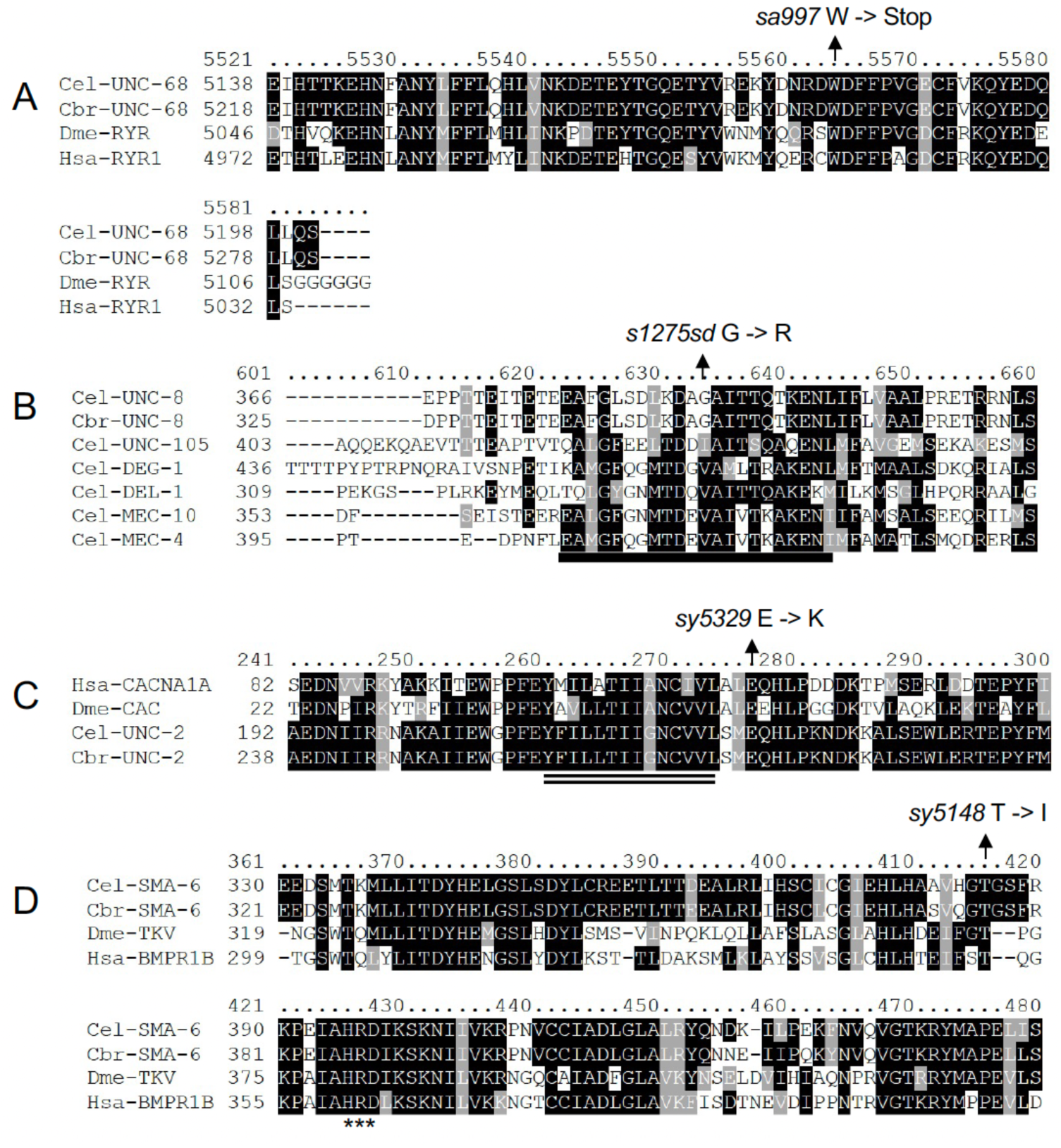
Amino acid sequence alignments for select proteins. A. A mutation in JT10275 results in a premature stop codon in *Cbr-unc-68* that truncates the predicted encoded protein, removing elements of a conserved carboxyl-terminus domain that functions in stabilization of the RYR channel pore and interacting with channel activating ligands (Peng et al., 2016; Clarke and Hendrickson, 2017). B. A mutation in BC1977 corresponds to a G to R substitution Cbr-UNC-8 in a domain common to DEG/ENaC family members shown to be important in channel closing (García-Añoveros et al. 1995). Alleles of *Cel-unc-8* (*e15*, *n491*) that confer a dominant phenotype similarly code for charged (G to E) substitutions impacting the same residue (Tavernarakis et al. 1997). The conserved DEG/ENaC domain is marked with a black line. C. The Unc phenotype in PS9334 coincides with a E to K substitution, affecting a conserved residue directly after the first transmembrane domain of the encoded calcium channel subunit alpha-1A protein. The predicted transmembrane domain is marked with a double line. D. The Dpy phenotype in PS9148 corresponds to a T to I substitution at a conserved residue within the kinase domain of Cbr-SMA-6. The HRD motif of the kinase domain catalytic loop is identified with three asterisks. Sequences aligned using Clustal Omega (Sievers et al. 2011) and shaded using boxshade (https://junli.netlify.app/apps/boxshade/). Identical amino acids are shaded with black, and conserved amino acids are shaded with gray. The sequences used in to produce the alignments are in Supplemental Table 15.

### Annotation of visible mutations linked to the physical map for an updated C. briggsae map resource

To place the ten genes mapped and characterized here in a larger genetic map context, we identified a set of 63 *C. briggsae* genes with mutant alleles reported in the literature (Figure 5, Supplemental Table 16). We included genes with alleles expected to disrupt or alter gene function and that were assigned to specific genes, either because the allele was generated using gene-targeting methods or because a mutation isolated based on phenotype was mapped and assigned to a gene. We then identified the physical position for each gene in the updated QX1410 genomic map (Supplemental Table 17). For this map, the relative distances represent physical, rather than genetic (recombination frequency) distances among the genes, because recombination frequency data are generally sparse and available for relatively few of the mutations. However, this improved physical map increases the confidence of the relative gene order and provides a resource of mutations available for future studies.

**Figure 5.**
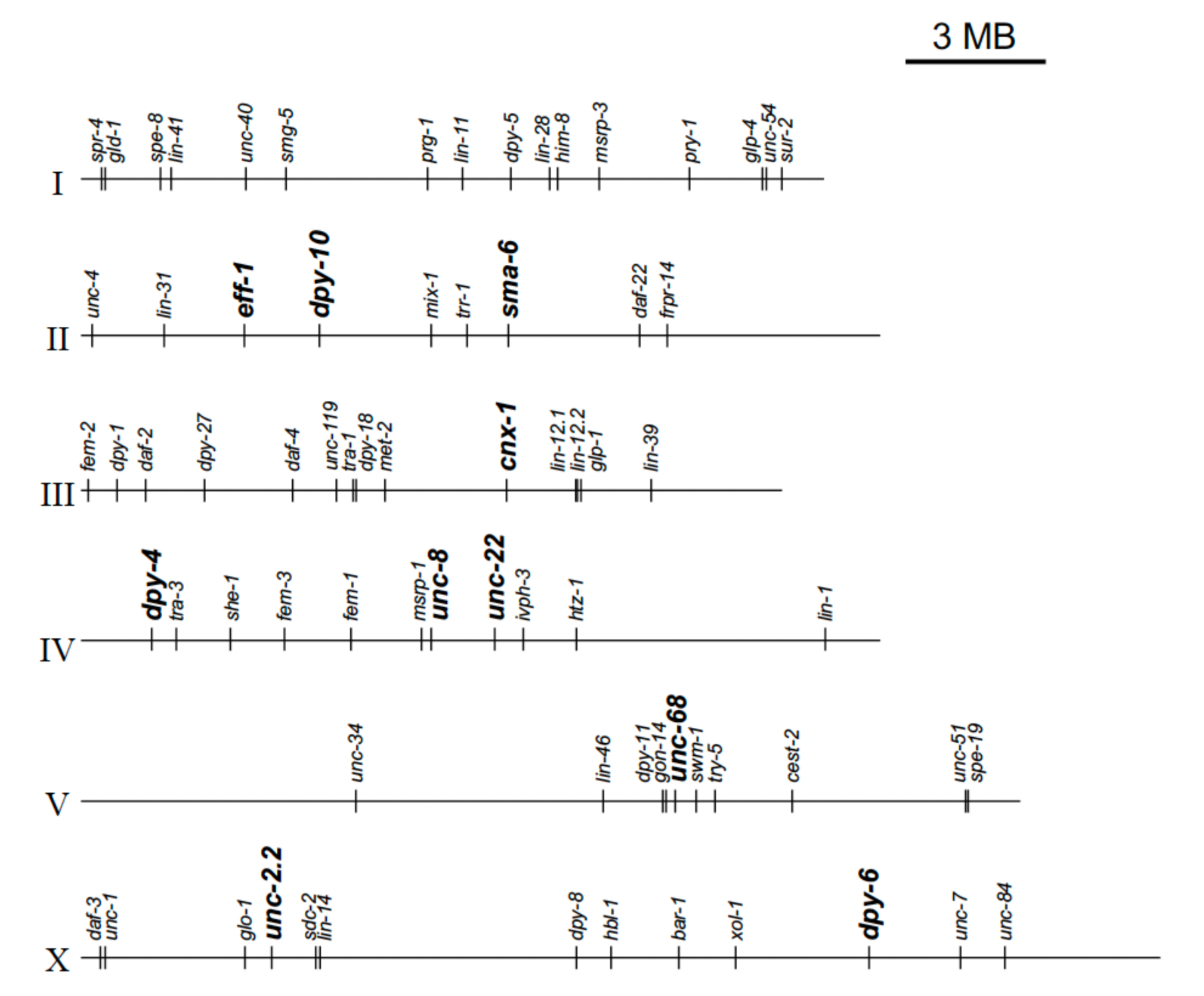
A map for the *C. briggsae* genome, including genes with mutations that are linked to the physical map. The physical location for *C. briggsae* genes with known mutations (Supplemental Table 17). The ten genes mapped in this project are indicated with larger font and in bold.

## Discussion

We identified, validated, and applied a set of indel variants with updated genomic sequence information to map genetic loci in the nematode *C. briggsae*. The primers used to identify each indel were evaluated against four *C. briggsae* wild strains, expanding their use to a range of comparative experiments and mutations isolated in different genetic backgrounds. Based on this set of validated indels and the potential for additional sites, VX34 represents a strong choice for mapping mutations isolated in the canonical wild-type background CGC2, due to the large number of indel differences between the strains. However, like HK104, regions of genomic incompatibility can skew the recovery of regional recombinants (Ross et al., 2011; Ben-David et al., 2021; Stevens et al., 2022). We demonstrated that variant mapping of *C. briggsae* mutations from a range of different labs produces sufficient genomic resolution that, in combination with whole-genome sequence information, we can identify prospective causative genes with high confidence despite the presence of many unique sequence variants in the background of each strain. Although we applied this approach to mutations with straight-forward visible mutant phenotypes, we consider this approach to be especially useful for traits that are difficult to reliably phenotype in the F2 offspring of an F1 heterozygote, including traits that exhibit maternal effects, variable expressivity, or incomplete penetrance. This approach is an important addition to the toolkit of methods available in which high-throughput sequencing is used as part of a pipeline for the identification of genes impacted in strains bearing novel mutations (Doitsidou et al. 2016).

This project mapped and identified genetic changes associated with ten genes, adding to the set of genes available for comparative genetic studies between *C. elegans* and *C. briggsae*. Three genes (*Cbr-dpy-4*/Collagen, *Cbr-dpy-6*/Mucin, *Cbr-dpy-10*/Collagen) are predicted to be involved in cuticle formation and two (*Cbr-unc-22*/Twitchin, *Cbr-unc-86*/Ryanodine Receptor) in muscle function. The alleles of the two genes anticipated to have neuronal functions (*Cbr-unc-8*/Na+channel, *Cbr-unc-2.2*/Ca++channel) each alter conserved residues, but they represent novel substitutions not previously studied in *C. elegans* (Mathews et al. 2003; Wang and Sieburth 2013). *Cel-cnx-1*/Calnexin encodes an ER chaperone protein important for ion channel biogenesis but also for stress response (Lee et al. 2005; Bai et al. 2018), and the *Cbr-cnx-1* mutant might be useful in comparative studies because of the differences between *C. elegans* and *C. briggsae* in response to some stressors (Kalinnikova et al. 2016; Jhaveri et al. 2025). *Cel-eff-1* mutants have been used to study cell fusion and to understand its roles in cell-fate specification, developmental progression, tissue homeostasis, morphogenesis, and wound healing, among other processes (Podbilewicz et al. 2006; Oren-Suissa et al. 2010; Pellegrino et al. 2011; Roiz et al. 2016; Ghose et al. 2018; Meng et al. 2020; Koneru et al. 2021; Funk et al. 2025). We anticipate that *Cbr-eff-1(s1281)* mutants will permit comparable studies in *C. briggsae*.

High-throughput sequencing and genome-editing methods have transformed comparative genomics and genetics, and these methods have been especially important in expanding the variety of organisms accessible to address research questions (e.g., Zhang et al. 2017; Karageorgi et al. 2019; Rasys et al. 2019; Shirai et al. 2022; Popsuj et al. 2024; Sommer-Trembo et al. 2024). Despite these advances, experiments using species of the *Caenorhabditis* genus retain several advantages due to their ease of culture and cryopreservation, rapid generation times, and relatively small genome sizes. In particular, these features make *Caenorhabditis* species, especially androdioecious ones such as *C. briggsae*, ideal for both targeted hypothesis-testing experiments but also large-scale forward genetic screens that have the capacity to identify novel or taxonomically restricted gene functions. The updated methods and resources in this report enhance *C. briggsae* as a platform for independent genetic discovery that is not anchored to orthology with *C. elegans*.

## Materials and methods

### Strains and maintenance

All nematode strains were grown on NGM plates seeded with *Escherichia coli* strain OP50 as a food source, following *C. elegans* protocols (Stiernagle 2006). Here, we use the strain name CGC2 to identify the standard reference *C. briggsae* strain and genome assembly (O’Connor et al. 2026). In some cases for clarity, we use CGC2 (AF16). Strains were grown and experiments performed at 20°C, unless otherwise noted. Specific strains and their genotypes are listed in Supplemental Table 1. For ease of reference, the strains with mapped and identified mutations are also listed here: BC1972 *Cbr-unc-22(s1270)* IV; BC1977 *Cbr-unc-8(s1275sd)* IV; BC1983 *Cbr-eff-1(s1281)* II; CM3378 *Cbr-dpy-4(sy5027)* IV; CP4 *Cbr-dpy-10(nm4)*; JT10275 *Cbr-unc-68(sa997)* V; PB101 *Cbr-dpy-6(bd101)* X; PS9148 *Cbr-sma-6(sy5148)* II; PS9334 *Cbr-cnx-1(sy5330)* III; *Cbr-unc-2.2(sy5329)* X.

### Genetic crossing methods for mapping

Males from the comparison strain (HK104 or VX34) were crossed with hermaphrodites from each mutant strain. F1 L4 hermaphrodites were plated individually and allowed to self-cross. F2 animals were selected for two different types of analysis. For bulk-segregant analysis, phenotypically mutant animals (at least 50) were selected to one (60 mm) plate and phenotypically wild-type animals (at least 50) were selected to a second plate. For individual F2 mapping, F2 animals exhibiting the mutant phenotype were selected and plated individually, except for the *s1275* allele, where phenotypically wild-type animals were selected instead of semi-dominant Unc. These plates were screened in the next generation to confirm that the trait breeds true. For both bulk segregant and individual F2 mapping, the plates were cultured until the animals cleared the bacteria (“starved plate”) and then processed for DNA recovery and PCR analysis.

### DNA preparation and PCR analysis

A small chunk of each starved plate was transferred to a fresh plate to preserve the line, and the remaining worms washed off using 1.5 ml M9 buffer and transferred to a 1.5 ml tube. Worms were collected by centrifugation for 1 minute, the supernatant removed with a pipette, and the tubes placed at −70°C overnight. 100 µl of Worm Lysis Buffer (50 mM KCl, 10 mM Tris (pH 8.2), 2.5 mM MgCl_2_, 0.45% NP-40, 0.45% Tween 20, 0.01% gelatin) plus proteinase K (at 0.1-0.5 µg/µl final concentration) was added to the frozen pellet, and the tubes incubated in a heat block at 65°C (with occasional vortexing) for 1 hour, followed by 95°C for 15 minutes to deactivate the proteinase K. 1.0 µl of this prepared DNA was used as template for PCR. In some cases, better results were obtained with 0.5 µl (or 1 µl of a 1 in 10 dilution) of this DNA.

All primers were evaluated with the same PCR conditions. Each 25 µl reaction contained 0.2 µl Taq and 1x buffer (final concentration; Syd labs), 200 µM (each) dNTPs, and 0.4 µM (each) primer. Reactions were completed on an MJ Research PTC thermocyler with initial denaturation at 94.0°C for 1 min; 35 cycles of 94.0°C 30 sec, 52.0°C 20 sec, 72.0°C 1 min; final extension at 72°C for 10 min. 10 µl of each sample was evaluated on a 2% agarose gel.

### DNA preparation and whole-genome sequencing

DNA extraction was performed following established protocols (Cook et al. 2016). In brief, to extract DNA, we transferred nematodes from three 10 cm NGMA plates spotted with OP50 *E. coli* into a 15 ml conical tube by washing with 10 mL of M9. We then used gravity to settle animals on the bottom of the conical tube, removed the supernatant, and added 10 mL of fresh M9. We repeated this wash method three times to serially dilute the E. coli in the M9 and allow the animals time to purge ingested *E. coli*. Genomic DNA was isolated from 100 to 300 µl nematode pellets using the DNAEasy Blood and Tissue isolation kit (cat# 69506, QIAGEN, Valencia, CA). The DNA concentration was determined for each sample with the Qubit dsDNA Broad Range Assay Kit (cat# Q32850 Invitrogen, Carlsbad, CA). Short-read libraries were generated with NEBNext® Ultra™ II FS DNA Library Prep (cat# E6177L) and Unique Dual Index primers from NEB (cat# E6440L or E6442). These libraries were sequenced on the Illumina NovaSeq 6000 (paired-end, 150 bp reads) at the New York University core facility or the Integrated Genomics Center at Johns Hopkins University.

### Sequence comparison to reference and mutation identification

Whole-genome short-read DNA sequencing from the six mutant strains reported here plus 13 other mutant strains and PB420 (now defined as CGC2) were aligned to the QX1410 reference genome using *GATK* (Van der Auwera et al. 2013) in a custom *Nextflow* (Di Tommaso et al. 2017) pipeline. Variants that were shared among the six mutant strains reported here plus the 13 additional strains (to be described in future studies) and CGC2 were classified as background variants and represent genetic differences between CGC2 and QX1410. The functional consequences of nuclear variants were predicted using *CSQ* from *bcftools* (Danecek and McCarthy 2017) and *VEP* (Hunt et al. 2022), and mitochondrial variants were annotated using *SnpEff* (Cingolani et al. 2012) using the invertebrate mitochondrial genetic code. Grantham (Grantham 1974) and BLOSUM62 (Henikoff and Henikoff 1992) scores were annotated using a custom script (https://github.com/AndersenLab/annotation-nf), in addition to the percent protein, or where in the protein a variant is present. Cytosine to thymine and guanine to adenine transitions were annotated as possible EMS-derived mutations.

To identify candidate gene for each strain, we used the breakpoints defined by the data representation of Supplemental Tables 8-14 to define an interval for each gene. Then using a hierarchical approach, we prioritized based on mutation (nonsense, frameshift, splicing, missense) and then on gene conservation (*C. elegans* ortholog vs CBG vs QX1410).

### Identification of published C. briggsae gene mutants

The search term “briggsae” was used to query the PubMed database. Relevant papers from 2006-June 2026 were reviewed, with possible gene mutations recovered from the Methods sections or supplemental strain lists. This process identified the 63 genes (Figure 5). These references are identified by PMID (Supplemental Table 16). Individual genes from this table were extracted (Supplemental Table 17) and supplemented with the genes identified as part of this work. The physical position for each gene was obtained from the genome browser on Caendr.org (release 20240129 (Crombie et al. 2024)), using the QX1410 assembly.

## Data Availability

Strains are available upon request. Sequence data are available publicly from NCBI accession PRJNA1551577. Other data necessary for confirming the conclusions of the article are present within the article, figures, and supplemental tables.

## Acknowledgements

We thank M. Ailion, D. Baillie, S. Baird, E. Haag, T. Inoue, P. Sternberg, J. Thomas and their research groups for generating mutations and providing strains. D. Baillie would like to acknowledge the National Research Council (NSERC) of Canada for their support of his research group. We enthusiastically thank the *Caenorhabditis* Genetics Center, which is funded by the NIH Office of Research Infrastructure Programs (P40 OD010440), for their essential role in curating, preserving, storing and finally distributing strains across both space and (especially in this case) time.

